# Bacteriophage T4 Vaccine Platform for Next-generation Influenza Vaccine Development

**DOI:** 10.1101/2021.06.14.448336

**Authors:** Mengling Li, Pengju Guo, Cen Chen, Helong Feng, Wanpo Zhang, Changqin Gu, Guoyuan Wen, Venigalla B. Rao, Pan Tao

## Abstract

Developing influenza vaccines that protect against a broad range of viruses is a public health priority, and several conserved viral proteins or domains have been identified as promising targets for such vaccine development. However, none of the targets is immunogenic, and vaccine platforms that can incorporate multiple antigens with enhanced immunogenicity are desperately needed. In this study, we provided proof-of-concept for the development of next-generation influenza vaccine using T4 phage virus-like particle (VLP) platform. With extracellular domain of influenza matrix protein 2 (M2e) as a readout, we showed that more than 1,280 M2e molecules can be assembled on a 120×90 nanometer phage capsid to form T4-M2e VLPs, which are highly immunogenic and induced complete protection against influenza virus challenge without any addition adjuvant. Potentially, additional conserved antigens or molecular adjuvants could be incorporated into the T4-M2e VLPs to customize influenza vaccines to address different issues. All the components of T4 VLP vaccines can be mass-produced in *E. coli* in a short time, therefore, providing a rapid approach to deal with the potential influenza pandemic.

## Introduction

Influenza A virus is a highly contagious agent that can cause severe respiratory diseases[1, 2]. Although vaccines are available, they are strain-specific and mainly target the variable head domain of viral major envelope glycoprotein, hemagglutinin (HA)[3, 4]. The stalk domain of HA exhibits some degree of conservation among influenza virus strains but cannot efficiently induce antibody responses in its native state due to the immunodominance of head domain[5, 6]. The rapid evolution of influenza viruses through antigenic drift and shift in their surface glycoproteins, HA in particular, makes current vaccines less efficient[7-10]. Therefore, vaccines have to be reformulated annually using reference viruses recommended by World Health Organization based on the information provided by their Global Influenza Surveillance Network[11].

Vaccines that provide broader protection against most influenza virus strains are highly desired. Many efforts have been focused on developing such universal vaccines using the conserved viral proteins or domains[12, 13]. Internal proteins nucleoprotein (NP) and matrix protein 1(M1) were mainly used as immunogens to induce cellular immune responses, particularly CD8+ T cells with cross-protection against heterologous influenza viruses[14-16]. Engineered headless HA stalk, in which the immunodominant head domain was removed, were mainly used to induce antibodies that recognize or neutralize diverse influenza virus strains[17, 18]. The extracellular of matrix protein 2 (M2e) is highly conserved among divergent influenza virus strains and was widely used as a target for universal vaccines[19, 20]. However, none of those vaccine targets are highly immunogenic, and many strategies were employed to enhance the immune responses[21, 22].

By taking advantage of *in vitro* assembly of antigen proteins on bacteriophage T4 capsid, we recently developed a virus-like particle (VLP) platform that can elicit robust immune responses against the displayed antigens without any adjuvants[23, 24]. In this study, we aimed to develop M2e influenza vaccines using the T4 VLP platform. The immunogenicity of M2e, a 23-residue peptide at the NH2-terminus of viral matrix protein 2 (M2), is quite low during natural infection due to its small size and low abundance on the virion surface[25]. However, when displayed on VLP, the M2e induced significant immune responses and provided variable protection against influenza virus infection[26, 27]. Although many different VLPs such as hepatitis B virus core particles [28], human papillomavirus particles [29], tobacco mosaic virus [30] were used as carriers for M2e, phage capsid particles are the most cost-effective and attractive ones due to their large-scale manufacturing potential, which could be critical during the influenza pandemic.

The M2e proteins displayed on T7 phage capsids are immunogenic but failed to provide completely protection against lethal influenza virus challenge, which probably because the low copy of M2e proteins per phage capsid [31]. Indeed, it was found that vaccines with higher M2e epitope densities resulted in higher protection efficacy [32, 33], and most of the licensed viral vaccines contain high density of antigens on the virion surface [34]. Although phage fd can display up to 2,700 copies of peptide per capsid through its major coat protein pVIII, the display is sensitive to the size of the peptide [35]. Therefore, only part of M2e (residues 2-16) was able to displayed on phage fd, which still provided complete protection but the challenged mice showed severe body weight loss [36]. We have showed that phage T4 can efficiently display full-length proteins as large as 120 kDa at high density because of its unique capsid architecture [37], which composed of four capsid proteins, gp23* (930 copies), gp24* (55 copies), Soc (small outer capsid protein, 870 copies), and Hoc (highly antigenic outer capsid protein, 155 copies) [23, 38]. Deletion of Hoc and Soc (Hoc^-^ Soc^-^ T4) has no effect on the propagation of T4 under lab conditions, and recombinant Hoc or Soc proteins can specifically bind to Hoc^-^Soc^-^ T4 capsids with high affinity [23, 24, 39, 40].

In current study, we showed that 3M2e proteins, which contain three tandem copies of M2e from human, swine, and avian influenza viruses can be efficiently displayed on Hoc^-^Soc^-^ T4 capsids with high density by fusion to COOH-terminal of Soc. The generated 3M2e-T4 VLPs were highly immunogenic and induced complete protection against lethal influenza virus challenge, without additional adjuvant. Most significant, the immunized mice show no or minor symptoms after influenza virus challenge based on clinical observations, body weight, and pathological analysis. Our studies provide proof-of-concept for the development of next-generation influenza vaccine using T4 VLP platform.

## Results

### Construction of the M2e-decorating T4 bacteriophages nanoparticles

To increase antigen density and the breadth of protection, a 3M2e gene containing three types of M2e from human, swine, and avian influenza virus was synthesized. The 3M2e gene was then fused to the COOH-terminus of Soc to generate Soc-3M2e, and three GGSSGGSS flexible linkers were used as indicated in Figure 1A to minimize interference between each component. Soc-3M2e fusion protein was expressed in *E. coli* and purified using nickel affinity chromatography followed by size exclusion chromatography, from which the major peak corresponding to a molecular weight of ∼21.8 kDa (monomeric Soc-3M2e) was collected (Fig. 1B, blue profile). The purity of the recombinant Soc-3M2e proteins was confirmed by sodium dodecyl sulfate–polyacrylamide gel electrophoresis (SDS-PAGE) showing the presence of a major band that is equivalent to the mass of Soc-3M2e protein (Fig. 1B).

**Figure 1.**
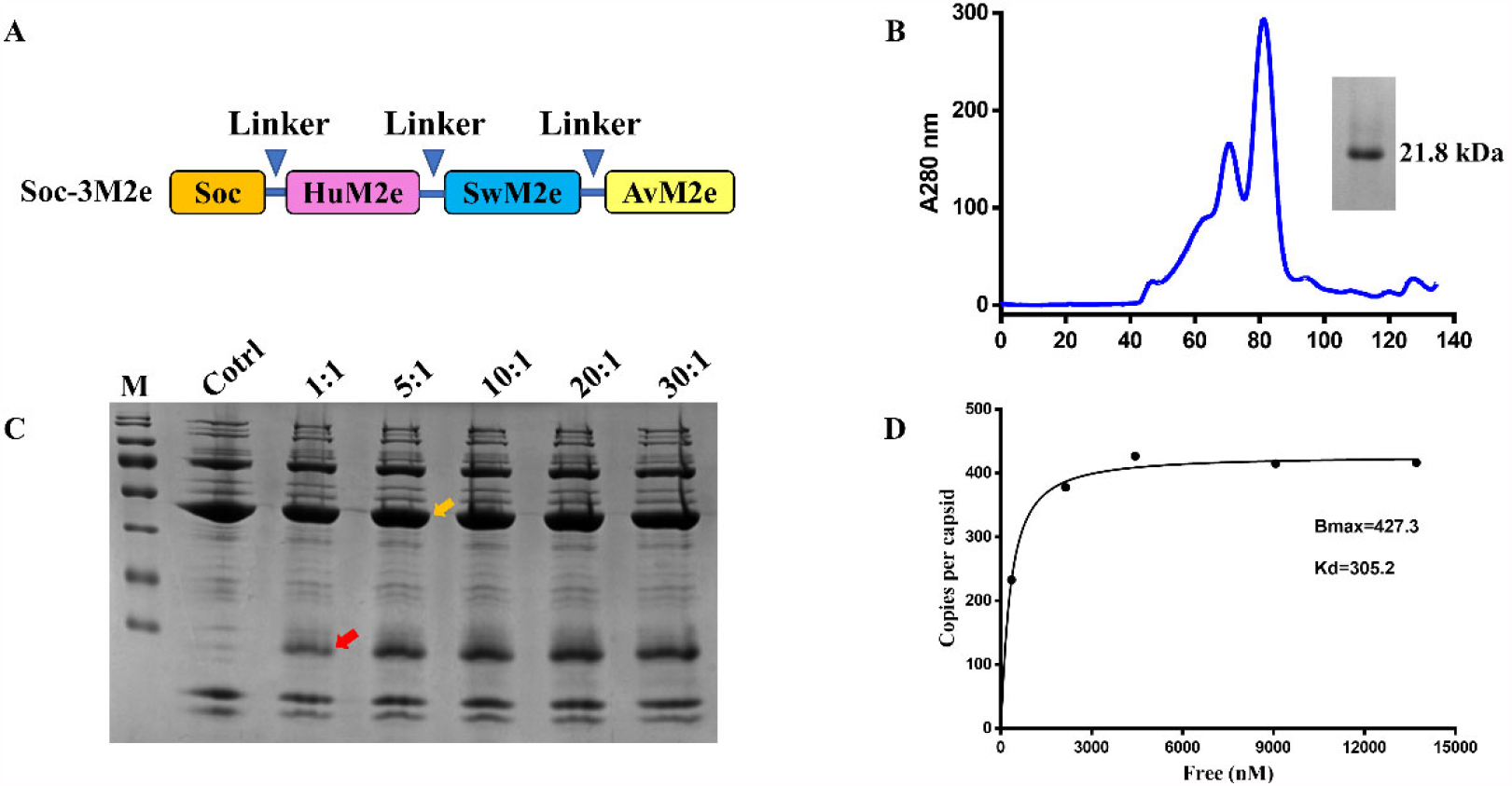
Construction of T4-3M2e VLPs. **(A)** Soc-3M2e fusion was constructed by fusing 3M2e, which contains three tandem copies of M2e from human, swine, and avian influenza viruses, to the COOH-terminus of Soc. Arrows indicated the flexible linkers (GGSSGGSS) between each component. Soc-3M2e protein was purified by HisTrap affinity chromatography followed by size exclusion chromatography **(B)**. Only the major peak was collected, and the purity of Soc-3M2e protein was analyzed by SDS-PAGE. **(C)** Assembly of T4-3M2e VLPs *in vitro*. About 5×10^10^ Hoc^-^ Soc^-^ T4 phages were incubated with at the indicated ratios of Soc-3M2e protein molecules to capsid binding sites (see Materials and Methods for the details). T4-3M2e VLPs were analyzed by SDS-PAGE. The same amount of Hoc^-^Soc^-^ T4 phages was used as a control. Yellow and black arrows indicated gp23* and Soc-3M2e, respectively. **(D)** Saturation binding curve of Soc-3M2e. The bound and unbound (not shown) Soc-3M2e proteins were calculated using BSA a standard. The copy numbers of Soc-3M2e per capsid were determined using gp23* as internal control. The data were plotted as one-site saturation ligand binding curve.

3M2e decorated T4 nanoparticles were prepared by incubation of Soc-3M2e proteins with Hoc^-^ Soc^-^ T4 phages as previously described [41]. To optimize the copy number of 3M2e, 5×10^10^ T4 phages were incubated with different amounts of Soc-3M2e proteins (Fig. 1C). The presence of 3M2e was determined by SDS-PAGE analysis of T4-3M2e nanoparticles. The Soc-3M2e proteins bound efficiently to the Hoc^-^Soc^-^ T4 phages even at a 1:1 ratio of Soc-3M2e molecules to Soc binding sites, and reached saturation at ratio of 10:1(Fig. 1C). The copy number of bound Soc-3M2e per capsid (Bmax) was 427.3, and the binding constants (Kd) were 305.2 nM. Since each Soc-3M2e protein contains three tandem copies of M2e peptide, there are more than 1,280 M2e molecules were assembled on each T4 nanoparticle, which is remarkably higher than any VLPs reported so far.

### T4-3M2e nanoparticles induced robust M2e-specific antibodies

To determine the immunogenicity of T4-3M2e, mice were intramuscularly immunized with T4-3M2e nanoparticles displaying 15μg 3M2e proteins on day 0, 14, and 28 (Fig. 2A). Mice immunized with PBS, 15μg of Soc-3M2e soluble proteins, or a mixture of 15μg Soc-3M2e proteins and phage T4 (3M2e+T4) were used as controls. To minimize the binding, Soc-3M2e proteins were mixed with phage T4 right before immunization. Sera were collected according to the scheme shown in Fig. 2A, and the titer of M2e-specific antibodies was determined by enzyme-linked immunosorbent assay (ELISA). All mice immunized with PBS were negative for M2e-specific IgG at the sera dilution of 50. The Soc-3M2e soluble proteins were able to induced low level of M2e-specific IgG antibodies, which was significant increased when Soc-3M2e proteins were mixed with T4 phages (Fig. 2B). This indicated the adjuvant activity of T4 phage nanoparticles. T4-3M2e nanoparticles, without additional adjuvant, induced highest level of 3M2e-specific IgG antibodies, up to end point titers of ∼4×10^5^ (Fig. 2B).

**Figure 2.**
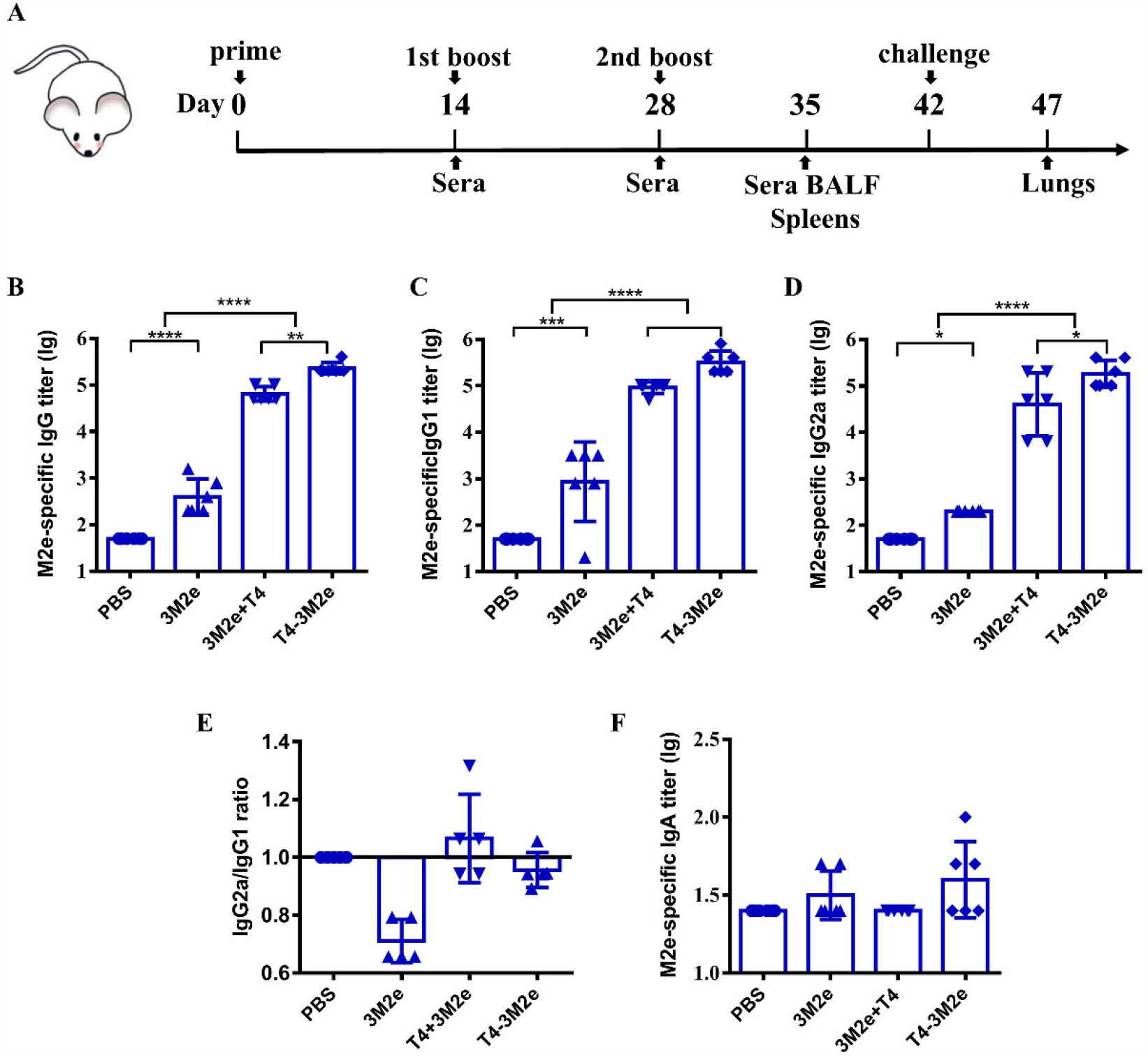
M2e-specific humoral immune responses. **(A)** Scheme of mouse immunization. Sera were obtained before each immunization. The titers of M2e-specific IgG **(B)**, IgG1**(C)**, IgG2a **(D)**, and IgA **(E)** were determined by ELISA using peptides pool of M2e from human, swine, and avian influenza viruses. **(F)** Ratio of M2e-specific IgG2a to IgG1 were calculated. Date was shown as means±S.D. *, **, and *** indicated p< 0.05, p< 0.01, and p< 0.001 respectively (ANOVA).

Since the M2e induced immune protection mainly depends on antibody-dependent cellular cytotoxicity (ADCC) and antibody-dependent cellular phagocytosis (ADCP), of which the efficiencies are different between IgG subtypes[42-44], we determined the titers of 3M2e-specific IgG1 and IgG2a. The Soc-3M2e soluble proteins mainly elicited IgG1, whereas mice immunized with T4-3M2e or a mixture of Soc-3M2e and phage T4 produced similar level of 3M2e-specific IgG1 and IgG2a antibodies (Fig. 2C-E). We also determined the 3M2e-specific IgA antibodies in sera. Although T4-3M2e induced relative higher level of IgA in some mice, the differences among all groups are not statistically significant (Fig.2F).

### T4-3M2e nanoparticles elicited strong cellular immune responses

To investigate M2e-specific cellular immune responses, mice were sacrificed 6 days after the second boost, and spleens were collected to isolate peripheral blood mononuclear cells (PBMCs). The number of INF-γ and IL-4 secreting cells were analyzed by ELISPOT using 10μg/ml M2e peptide as a stimulus. Mice immunized with Soc-3M2e soluble proteins were incapable to generate INF-γ and IL-4 secreting cells, whereas mice immunized with T4-3M2e developed significantly more M2e-specific INF-γ (Fig.3A) and IL-4 secreting cells (Fig.3B). However, the mixture of Soc-3M2e and phage T4 cannot stimulate mice to develop M2e-specific INF-γ and IL-4 secreting cells (Fig.3), indicating that assembly of M2e nanoparticles is necessary for eliciting cellular immune responses.

**Figure 3.**
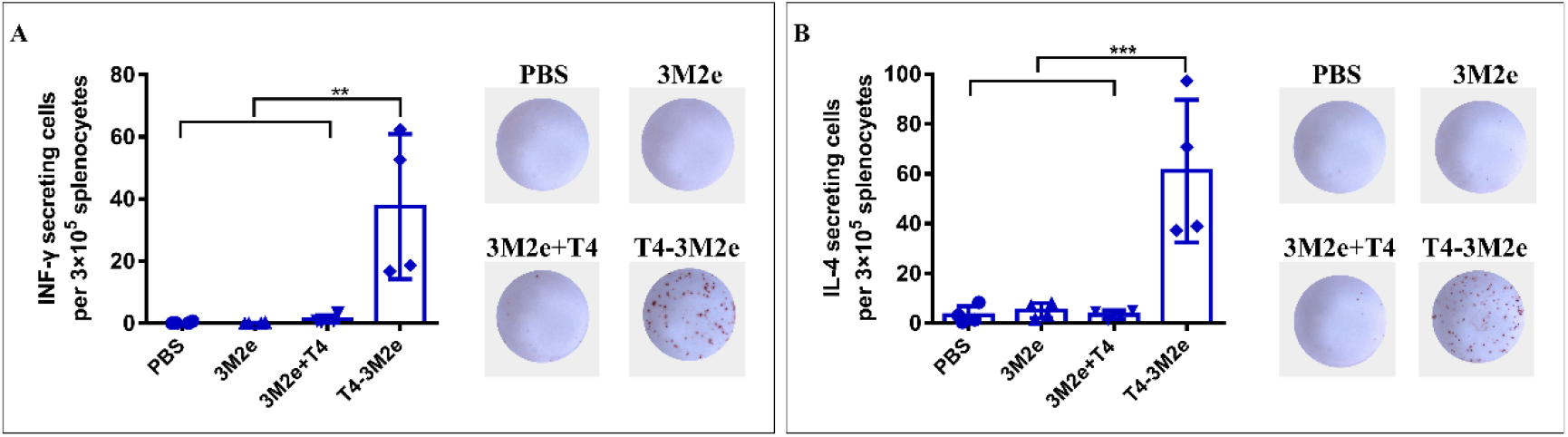
M2e-specific cellular immune responses. Mice (n=4) were immunized according to the scheme shown in Fig. 2A, and splenocytes were isolated on day 34. The INF-γ **(A)** and IL-4 **(B)** secreting lymphocytes were assayed by ELISPOT as described in Materials and Methods. The right images of each panel show the representative results of ELISPOT wells from each group. Data were represented as mean ± S.D. of four mice in each group. ** p < 0.01; *** p < 0.001 (ANOVA).

### T4-3M2e nanoparticles elicited M2e-specific mucosal antibodies

The mucosal surfaces of the respiratory tract are major ports of entry for influenza viruses, and previous studies indicated that both mucosal IgG and serum IgG are conducive to the defense response[45]. To determine the M2e-specific mucosal antibodies, bronchoalveolar lavage fluid (BALF) of mice was collected 7 days post last immunization, and the presence of IgG and IgA antibodies were detected using ELISA. As shown in Fig. 4A, soluble Soc-3M2e protein was able to induce low level of M2e-specific IgG antibodies, which can be significantly increased when the Soc-3M2e proteins were mixed with T4 phages. The 3M2e proteins assembled on T4 phages (T4-3M2e VLPs) induced highest level of M2e-specific IgG antibodies. As expected, the PBS control mice were negative for anti-M2e IgG antibodies (Fig. 4A). Analysis of M2e-specific IgG antibody subtypes showed that Soc-3M2e soluble proteins only elicited IgG1, whereas mice immunized with T4-3M2e or a mixture of Soc-3M2e and phage T4 induced similar level of 3M2e-specific IgG1 and IgG2a antibodies (Fig. 4B and C). However, all groups were failed to developed M2e-specific IgA antibodies in BALF (Fig. 4D).

**Figure 4.**
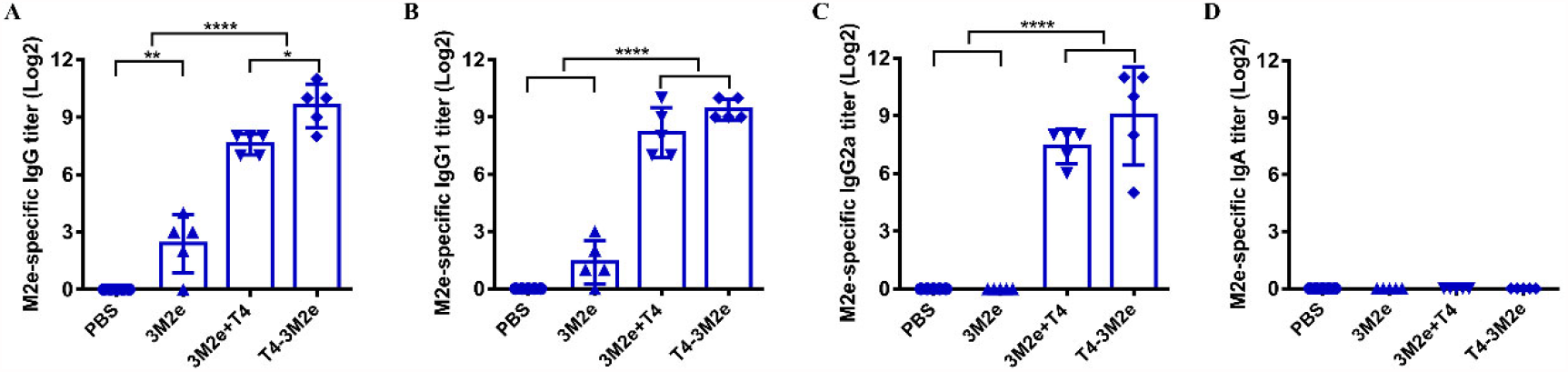
M2e-specific antibodies in BALF. Bronchoalveolar lavage fluids were collected 7 days after last immunization (n=5). M2e-specific total IgG **(B)**, IgG1**(C)**, IgG2a **(D)**, and IgA **(E)** were determined by ELISA using a mixture of human, swine, avian influenza virus M2e peptides as the capture antigen (2 µg/ml). Date was presented as means±S.D. *, p<0.05; **, p<0.05; ***, p<0.001 (ANOVA)

### T4-3M2e nanoparticles provided complete protection against influenza A virus challenge

To evaluate the protective efficacy of each formulation, immunized mice were challenged with 5LD_50_ of A/Puerto Rico/8/34 (H1N1) virus and monitored daily for survival and body weight for 14 days. As shown in Figure 5A, infection of influenza virus resulted in significant weight loss of mice immunized with PBS, Soc-3M2e soluble protein, or T4+3M2e three days post infection. All mice in PBS group died 10 days post the challenge. One of six mice immunized with soluble Soc-3M2e recovered from body weight loss and was survival, whereas 67% of mice vaccinated with T4+3M2e was survival (Fig. 5B). All mice immunized with T4-3M2e nanoparticles survived the lethal challenges with H1N1 virus infections, particularly, none of them showed obviously body weight loss (Fig. 5A and B). The protection efficacy was further evaluated by pathological analysis of lungs of immunized mice five days post-challenge with 5LD_50_ of A/Puerto Rico/8/34(H1N1). Figure 5C showed the representative results of overall picture of lung lesions (column I) and pathological changes of alveoli (column II), bronchi (column III), and pulmonary vessels (column IV). Overall, mice immunized with T4+3M2e mixture showed obvious but less severe lesions in the lungs than that of mice immunized with PBS or Soc-3M2e, whereas no obvious lesions were found in T4-3M2e immunized mice (column I). The alveolar wall of mice immunized with PBS or Soc-3M2e protein was severely thickened (column II), and a large number of inflammatory cells infiltrated around the bronchi (column III) and pulmonary blood vessels (column IV). Mice immunized with T4+3M2e mixture also exhibited similar but less severe pathological changes. However, mice vaccinated with T4-3M2e nanoparticles showed relatively normal alveolar wall thickness and neglectable inflammatory infiltration (Fig. 5C).

**Figure 5.**
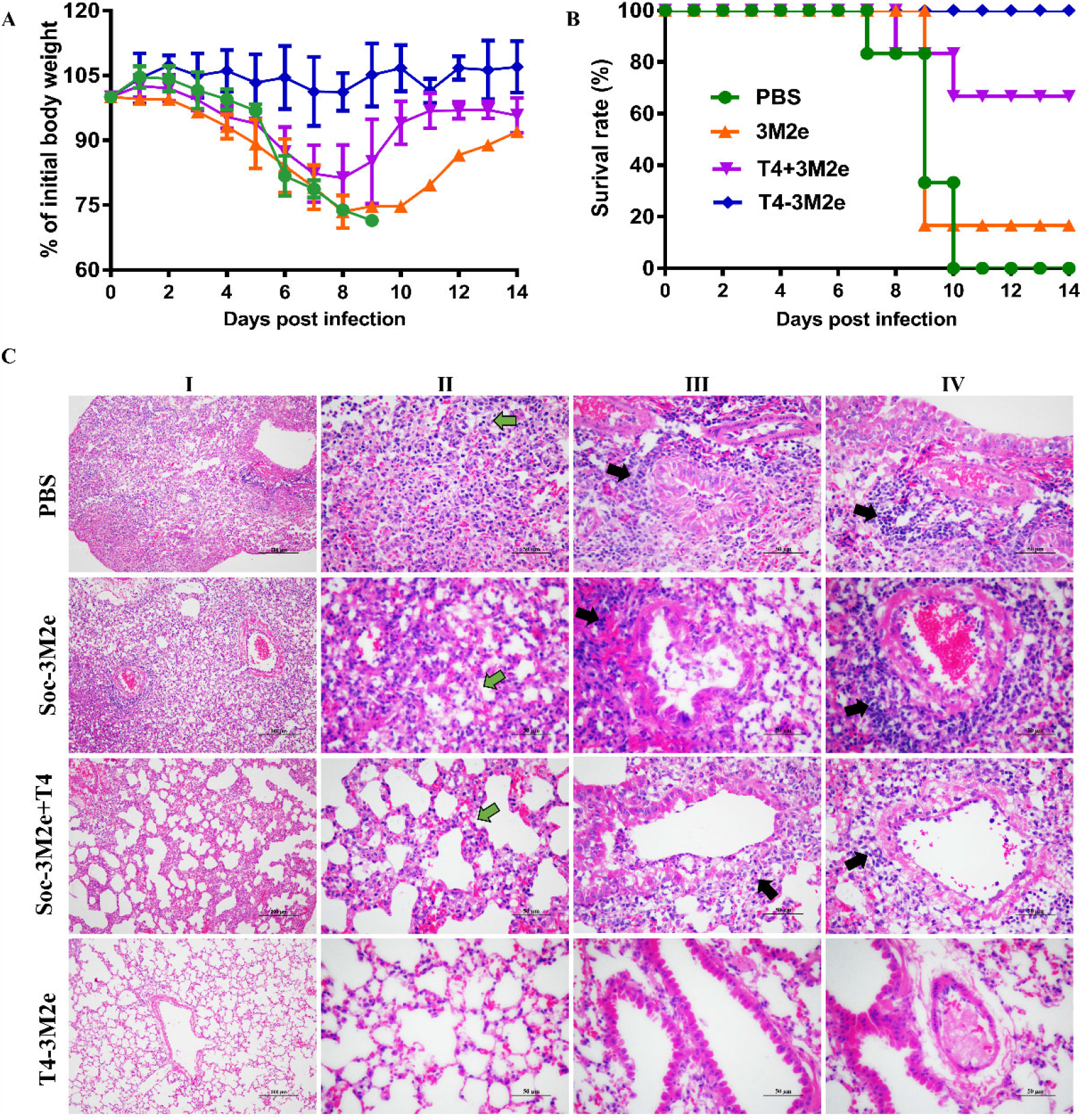
T4-3M2e VLPs provide complete protection against influenza virus challenge. Mice (n=6) were challenged with 5×LD_50_ of A/PR/8/34 two weeks after last immunization. Weight loss **(A)** and survival rate **(B)** of mice were monitored daily for 14 days. **(C)** Pathological analysis of lungs from mice (n=3) challenged with virus was carried out in a separate experiment, in which mice were immunized with the same immunization procedure described above. Five days post infection with 5×LD_50_ of A/PR/8/34, mice were euthanized, and lung sections were prepared as described in Materials and Methods. The representative results from each group were shown (column I, scale bar, 200 µm; Columns II-IV, scale bar, 200 µm). Main pathological changes are thickening of alveolar septa of mice (column II, green arrows), inflammatory cells infiltrated around the bronchi (column III), and pulmonary blood vessels (column IV). Black arrows indicate the inflammatory cell infiltration around the bronchi and pulmonary blood vessels.

## Discussion

M2e has been considered an attractive target for the development of universal influenza vaccines [46-48]. However, carriers and adjuvants are absolutely needed for M2e vaccines because of the poor immunogenicity of M2e peptides. By taking advantage of phage T4 platform, in this study, we developed a novel M2e VLP vaccine that, without additional adjuvant, induced robust immune responses and provided complete protection against influenza virus challenge.

The T4-M2e VLP vaccine was prepared by simply incubating Soc-3M2e fusion proteins with Hoc^-^Soc^-^ T4 phages, both of which can be produced in *E. coli* in large-scale. Therefore, our T4-M2e VLPs provide an approach to manufacture influenza vaccines with unlimited doses in a short time, which is critical to deal with emerging influenza pandemics. Although phages T7 and fd have also been used as carriers to present M2e, they induced limited protections [31, 49]. This probably because they cannot present full-length M2e in high density, which is the key determinant of immune responses induced by M2e-based vaccines [50]. We showed that ∼427 copies of the 21.8 kDa Soc-3M2e containing three tandem copies of M2e from human, swine, and avian influenza viruses were assembled on each T4 capsid. This means that 1302 copies of M2e molecules were presented on a 90×120 nanometer particle, which is higher than any VLPs reported so far. The 3M2e-T4 VLPs were highly immunogenic and induced robust immune responses without any adjuvants.

Apart from high epitope density, the high immunogenicity of T4-M2e VLP also probably because T4 phage has the ability to stimulate innate immune responses and act as a natural adjuvant [23, 35]. Indeed, we found the mixture of Soc-3M2e and T4 phages (most of the Soc-3M2e proteins did not bind to capsid and existed mainly as soluble proteins, see Results for the details) induced higher level of M2e-specific antibodies in sera (Fig. 2B-D) and BALF (Fig. 4). Significantly, T4-M2e VLP elicited even much stronger immune responses than that of the mixture of Soc-3M2e and T4 phages. This might because display of Soc-3M2e protein on T4 phage links the antigen to an adjuvant-loaded delivery system, which ensures simultaneous delivery of them to the same immune cell such as the antigen-presenting cells (APCs) that could significantly enhance the immune responses [51].

Unlike soluble Soc-3M2e protein that mainly elicit Th2-biased responses, we found T4-M2e VLPs induced balanced Th1 and Th2 immune responses (Fig.2F and Fig.3), which is vital for vaccines. Th1-type cytokines such as IFN-γ tend to induce the proinflammatory responses, which will result in tissue damage if beyond control. Th2-type cytokines such as IL-4 are mostly involved in mediating anti-inflammatory response, which will counteract the excessive microbicidal effect mediated by Th1-based responses [52-54]. Our results showed that the mice immunized with T4-M2e VLPs, but not soluble Soc-3M2e proteins, produced similar level of M2e-specific IgG1 and IgG2a (Fig.2F). Similar results were also observed for the M2e-specific INF-γ and IL-4 secreting cells (Fig.3) indicating that T4-M2e VLPs facilitate both Th1-type and Th2-type immune responses. Additionally, M2e-specific antibodies generally show non-neutralizing activity and their protections mainly depend on antibody-dependent cellular cytotoxicity (ADCC) and antibody-dependent cellular phagocytosis (ADCP), which assist in viral clearance [42-44, 55]. Therefore, high level of M2e-specific IgG2a antibodies, which are more potent than other IgG subclasses in directing ADCC, is desirable for M2e-based vaccines.

Previous studies have suggested that T-cell responses induced by M2e vaccines also contributed to the protection against influenza infections[47, 56]. In our current study, we found T4-M2e VLPs, rather than the mixture of T4 and Soc-3M2e, elicited high level of M2e-specific INF-γ and IL-4 secreting lymphocytes in the spleens of vaccinated mice, which most likely related to the cross-presentation of particle antigens. This might contribute to the enhanced protections of T4-M2e VLPs as shown in Figure 5.

Although M2e-based universal influenza vaccines are promising, it is highly desirable to include other conserved antigens such as HA-stalk, NP, and M1 to cover a broad range of virus types. Other than display of M2e on capsids, our T4 vaccine platform provides huge room for next-generation vaccine design. For instance, we have demonstrated T4 platform can be used to simultaneously display and deliver different kinds of protein antigens, deliver antigens proteins or DNAs encoding antigen proteins, specifically deliver antigens to dendritic cells, and co-deliver antigens and molecular adjuvants[24, 35, 37]. Therefore, more antigens can be incorporated into T4-M2e VLPs either as proteins or DNAs, and generated VLP vaccines can be targeted to dendritic cells by display of targeting molecules. Such related experiments are underway to develop next-generation influenza vaccines.

In conclusion, T4 phage platform can display full-length of 3M2e proteins with high density to generated T4-M2e VLP vaccine, which induced both strong humoral and cellular immune responses. The T4-M2e VLP vaccine, without additional adjuvant, completely protect immunized mice against lethal challenges of influenza virus. Our studies provide proof-of-concept for the development of next-generation influenza vaccine using T4 VLP platform. Potentially, additional conserved influenza antigens could be incorporated either as proteins or DNAs. In addition, the immunogenicity of T4 VLP vaccines could be further enhanced by targeting to immune cells or incorporating molecular adjuvants.

## Materials and Methods

### Ethics statement

This study was carried out in the Laboratory Animal Center of Huazhong Agricultural University. All mouse studies were approved by the Research Ethics Committee (HZAUMO-2021-0023), Huazhong Agricultural University, Hubei, China and performed strictly in accordance with the Guidelines for the Care and Use of Laboratory Animals that was approved by the Research Ethics Committee, Huazhong Agricultural University, Hubei, China.

### Design, construction, and preparation of recombinant 3M2e protein

Three conserved sequences of M2e extracellular domain were designed from human, swine, and avian influenza A viruses. The flexible linker (GGSSGGSS) sequences were inserted among three different M2e sequences, and the designed tandem M2e sequence was synthesized commercially after codon optimization of the sequences. Subsequently, M2e sequences were used to subclone into expression vector. The sequences are listed in the Supporting Information table 1.

For expression of Soc-3M2e fusion protein. The synthesized DNA was cloned into pET28b expression vector that has inserted into RB69 Soc gene, which resulted in 3M2e gene was fused to the C-terminal of RB69 Soc gene. The Soc-3M2e protein expression plasmids were transformed into Escherichia coli BL21 (DE3) competent cells. A single clone of transformed strains was cultured in LB medium supplemented with 50 mg/L of kanamycin overnight at 37°C, which was transferred to fresh LB medium at a ratio of 1:100. The bacteria were induced with 1mM IPTG at 30°C for 2h when the OD600 reach 0.6-0.8 following bacteria were collected by centrifugation at 7,000 rpm for 15min. Subsequently, bacteria were resuspended in binding buffer (20 mM Tris-HCl, 100 mM NaCl, 10mMimidazole, pH 8.0) supplemented with 5 μg/ml DNase I and broken by high-pressure cell disruptor at 4°C. Cell debris was removed by high-speed centrifugation (35,000g, 20 min, 4°C), and the supernatant passed through a 0.22 μm filter for protein purification preparation.

For protein purification, filtered supernatants was loaded onto Ni-NTA-resin equilibrated with binding buffer. The resin was washed with wash buffer (20 mM Tris-HCl, 100 mM NaCl, 20 mM imidazole, pH 8.0) and eluted with elution buffer (20 mM Tris-HCl, 100 mM NaCl, 400 mM imidazole, pH 8.0) according to the instructions. The concentration of collected protein in each tube was determined used by Bradford assay, and then 5 ml of the peak protein was forced through 0.22 μm filter. The proteins were further purified by size exclusion chromatography (Hi-load 16/60 Superdex 200 column, GE Healthcare Life Sciences) in a buffer (20 mM Tris-HCl, 100 mM NaCl, pH 8.0) performed on a purifier system. The proteins were further characterized by SDS-PAGE, and protein concentrations were determined by BCA assay.

### Assembled 3M2e antigens on the T4 capsid in vitro

The propagation and purification of Hoc^-^Soc^-^ phage T4 as previously described[41, 57]. Overnight-cultured E. coli strains P301 was inoculated to mixed LB and M9CA medium. when the density of E. coli strains P301 reach a 2×10^8^/ml concentration, E. coli was infected with Hoc^-^Soc^-^ phage T4 at a multiplicity of infection (MOI) of 0.2-0.4, which were allowed to culture at 37 °C until abundant cell debris was produced. Following bacteria were further lysed with chloroform at 37 °C, 180 rpm for 20 min, the mixture was collected by centrifugation at 30,000 g for 30 min. Mixture was suspended in Pi-Mg buffer (26 mM Na_2_HPO_4_, 22 mM KH_2_PO_4_, 79 mM NaCl, 1 mM MgSO_4_), and bacteria cells were further removed by centrifugation at 4,300 g for 20 min, followed incubated with chloroform and DNase I at 37°C, 180 rpm for 20 min. Then, the sediments were discarded and supernatant was centrifuged at high speed (30,000 g, 30 min) and samples were resuspended in 1 ml Pi-Mg buffer. The phage particles were purified by CsCl step density gradient centrifugation. Finally, the phages particles were dialyzed at dialysis solution I (10 mM Tris, 200 mM NaCl, 5 mM MgCl_2_, pH 8.0) for 5h and dialysis solution II (10mM Tris, 50mM NaCl, 5mM MgCl_2_, pH 8.0) overnight at 4 °C.

For *in vitro* assembly, about 5.6×10^10^ phage particles were centrifuged at 21,130 g (4 °C) for 45 min. After two times washes with phosphate buffered saline (PBS), the phage pellet was resuspended in PBS. An equal number of Hoc^-^Soc^-^ phage T4 phage particles were incubated with increasing amounts of Soc-3M2e at 4 °C for 45 min. The unbound Soc-3M2e proteins were removed by centrifugation at 21,130 g for 30 min, and the phage particles containing bound proteins were washed twice with PBS. The final phage particles were resuspended in PBS and transferred to a new tube.

The phage particles containing bound proteins was analyzed by SDS-PAGE, and Soc-3M2e proteins were quantified by Image-Pro Plus software. Quantify of gp23* and Soc-3M2e protein in each lane separately was calculated with different gradient BSA protein as standard. The copy number of bound Soc-3M2e proteins was determined using gp23 as the internal control which has a molecular mass of 49 kDa and expressed 930 copies per capsid has been identified. A saturation binding curve was formed as previously described[23, 24, 39]. Briefly, the copy number at the different ratios was shown on the y-axis, while the concentration of unbound antigen in each ratio was shown on the x-axis. Saturation binding curve is formed using Prism GraphPad software, and the binding Kd and the maximal number of binding sites (Bmax) were determined by nonlinear regression analysis.

### Immunizations and influenza A virus challenge

Female BALB/c mice (6-8 weeks) administered intramuscularly at week 0, 2, and 4. Mice vaccinated with PBS, 15μg of 3M2e soluble proteins, a mixture of 15μg 3M2e proteins and Bacteriophage T4, and T4-3M2e nanoparticles (15μg 3M2e proteins). All immunized mice besides control group received the same number of phage T4 particles.

Two weeks post the last immunization, mice (n=9) were intranasally infected with the lethal dose(5LD_50_) of influenza A/Puerto Rico/8/1934 (PR8) virus under anaesthetized with ether. All mice were monitored daily survival and weight loss for14 days. If the animal with above 30% weight loss, mice are euthanized immediately.

### Antibody detection in sera and BALF

The levels of antigen-specific antibodies in sera and BALF were determined by enzyme-linked immunosorbent Assay (ELISA). Briefly, 7 days after the last immunizations, sera and bronchoalveolar lavage fluid (BALF) were collected from mice. Mice blood was obtained from tail vein, and BALF was collected by the following treatment: after tracheal puncture, BALF was collected by flushing lung several times with 1ml PBS. The supernatant was subsequently harvested by centrifugation and stored at -20 °C until analysis.

After sampling collection, 96-well plates coated overnight at 4°C with 2 μg/ml mixing M2e peptides (consist of equal peptides from human M2e, swine M2e and Avian M2e) in 100 μl sodium bicarbonate buffer. The plates were blocked with 200 μl/well of PBS containing 3% BSA and 0.05% Tween-20 for 1 hour at 37 °C. Sera and BALF were serially diluted in PBS supplemented with 1% BSA and 0.05% Tween-20 and added to each well (100 μl/well), which incubated at 37°C for 1 hour. Then, HRP-conjugated goat anti-mouse IgG, IgG1, IgG2a, and IgA secondary antibodies diluted in PBS were added (100 μl/wells) and further incubated at 37 °C for 1 hour. 100 μl TMB as substrate was added to each well and then color development was stopped with 50 μl 2 M H_2_SO4. Finally, absorbance of OD450 was determined by a microplate reader. The ELISA endpoint titers were defined as value of sample dilution was twice the average value of the background.

### Cytokine ELISPOT

ELISpot assay was performed to determine the number of INF-γ and IL-4 secreting cells in spleen according to the manufacturer’s protocol (DAKAWE, China). Briefly, spleens were harvested from mouse at 7 days post the last immunizations and processed into single-cell suspensions. single-cells suspensions were obtained by mechanically dissociated in 4 ml lymphocyte isolation fluid, and filter the tissues through 70μm cell strainers. 3×10^5^ splenocytes were added to each well of plate and stimulated with M2e peptides (consist of equal amount of peptides from human M2e, swine M2e and Avian M2e) at a final concentration of 10 μg/ml. After cells cultured at 37 °C and 5% CO2 for 32-34 hours, splenocytes were removed by cell-cracking buffer of ice-cold deionized water. Then, plates were incubated with biotinylated antibody and followed treated with diluted HRP-conjugated streptavidin. Finally, the reaction was developed with AEC substrate and stopped with flowing water. Plates was dry naturally at room temperature, and spot counting was performed by the company.

### Histology assay

Five days post challenged with 5LD50 of influenza A/Puerto Rico/8/1934 (PR8) virus, lungs tissues were isolated from immunized mice (n=3). Lung tissues were fixed in 10% formalin and dehydrated through a graded series of ethanols. Then, tissues were embedded in paraffin wax, and each sample were cut at a thickness of 4 µm. Finally, deparaffinized sections from lungs were stained with hematoxylin-eosin.

### Statistical Analysis

All the statistical analysis in this study were performed using GraphPad Prism software. Comparisons among different groups were evaluated by one-way ANOVA. In all cases, p< 0.05 was considered as statistically significant difference.

## Acknowledgements

This work was supported by grants from National Natural Science Foundation of China [Grant No. 31870915 to PT], Fundamental Research Funds for the Central Universities [Program No. 2662019PY002 to PT], and in part by National Institute of Allergy and Infectious Diseases, National Institutes of Health [AI081726 to VBR], and National Science Foundation [MCB-0923873 to VBR].

## Reference

1. Monto, A.S., and Fukuda, K. (2020). Lessons From Influenza Pandemics of the Last 100 Years. Clinical infectious diseases : an official publication of the Infectious Diseases Society of America 70, 951–957.

2. Elbahesh, H., Saletti, G., Gerlach, T., and Rimmelzwaan, G.F. (2019). Broadly protective influenza vaccines: design and production platforms. Current opinion in virology 34, 1–9.

3. Jang, Y.H., and Seong, B.L. (2019). The Quest for a Truly Universal Influenza Vaccine. Frontiers in cellular and infection microbiology 9, 344.

4. Sautto, G.A., Kirchenbaum, G.A., and Ross, T.M. (2018). Towards a universal influenza vaccine: different approaches for one goal. Virol J 15, 17.

5. Wong, S.S., and Webby, R.J. (2013). Traditional and new influenza vaccines. Clinical microbiology reviews 26, 476–492.

6. Jang, Y.H., and Seong, B.L. (2014). Options and obstacles for designing a universal influenza vaccine. Viruses 6, 3159–3180.

7. Webster, R.G., and Govorkova, E.A. (2014). Continuing challenges in influenza. Annals of the New York Academy of Sciences 1323, 115–139.

8. Wei, C.J., Crank, M.C., Shiver, J., Graham, B.S., Mascola, J.R., and Nabel, G.J. (2020). Nextgeneration influenza vaccines: opportunities and challenges. Nature reviews. Drug discovery 19, 239–252.

9. Sah, P., Alfaro-Murillo, J.A., Fitzpatrick, M.C., Neuzil, K.M., Meyers, L.A., Singer, B.H., and Galvani, A.P. (2019). Future epidemiological and economic impacts of universal influenza vaccines. Proceedings of the National Academy of Sciences of the United States of America 116, 20786–20792.

10. Jazayeri, S.D., and Poh, C.L. (2019). Development of Universal Influenza Vaccines Targeting Conserved Viral Proteins. Vaccines 7.

11. Pica, N., and Palese, P. (2013). Toward a universal influenza virus vaccine: prospects and challenges. Annual review of medicine 64, 189–202.

12. R, N., and medicine, P.P.J.A.r.o. (2020). Is a Universal Influenza Virus Vaccine Possible? 71, 315–327.

13. Wei, C.J., Crank, M.C., Shiver, J., Graham, B.S., Mascola, J.R., and Nabel, G.J. (2020). Nextgeneration influenza vaccines: opportunities and challenges. Nature reviews. Drug discovery.

14. Li, Z.T., Zarnitsyna, V.I., Lowen, A.C., Weissman, D., Koelle, K., Kohlmeier, J.E., and Antia, R. (2019). Why Are CD8 T Cell Epitopes of Human Influenza A Virus Conserved? Journal of virology 93.

15. Herrera-Rodriguez, J., Meijerhof, T., Niesters, H.G., Stjernholm, G., Hovden, A.O., Sorensen, B., Okvist, M., Sommerfelt, M.A., and Huckriede, A. (2018). A novel peptide-based vaccine candidate with protective efficacy against influenza A in a mouse model. Virology 515, 21–28.

16. Pleguezuelos, O., James, E., Fernandez, A., Lopes, V., Rosas, L.A., Cervantes-Medina, A., Cleath, J., Edwards, K., Neitzey, D., Gu, W., et al. (2020). Efficacy of FLU-v, a broadspectrum influenza vaccine, in a randomized phase IIb human influenza challenge study. NPJ vaccines 5, 22.

17. Li, C.K., Rappuoli, R., and Xu, X.N. (2013). Correlates of protection against influenza infection in humans--on the path to a universal vaccine? Current opinion in immunology 25, 470–476.

18. Krammer, F. (2015). The Quest for a Universal Flu Vaccine: Headless HA 2.0. Cell host & microbe 18, 395–397.

19. Zharikova, D., Mozdzanowska, K., Feng, J., Zhang, M., and Gerhard, W. (2005). Influenza type A virus escape mutants emerge in vivo in the presence of antibodies to the ectodomain of matrix protein 2. Journal of virology 79, 6644–6654.

20. Nachbagauer, R., and Palese, P. (2020). Is a Universal Influenza Virus Vaccine Possible? Annual review of medicine 71, 315–327.

21. He, F., Leyrer, S., and Kwang, J. (2016). Strategies towards universal pandemic influenza vaccines. Expert Rev Vaccines 15, 215–225.

22. Deng, L., Mohan, T., Chang, T.Z., Gonzalez, G.X., Wang, Y., Kwon, Y.M., Kang, S.M., Compans, R.W., Champion, J.A., and Wang, B.Z. (2018). Double-layered protein nanoparticles induce broad protection against divergent influenza A viruses. Nature communications 9, 359.

23. Tao, P., Mahalingam, M., Kirtley, M.L., van Lier, C.J., Sha, J., Yeager, L.A., Chopra, A.K., and Rao, V.B. (2013). Mutated and bacteriophage T4 nanoparticle arrayed F1-V immunogens from Yersinia pestis as next generation plague vaccines. PLoS pathogens 9, e1003495.

24. Tao, P., Mahalingam, M., Zhu, J., Moayeri, M., Sha, J., Lawrence, W.S., Leppla, S.H., Chopra, A.K., and Rao, V.B. (2018). A Bacteriophage T4 Nanoparticle-Based Dual Vaccine against Anthrax and Plague. mBio 9.

25. Schepens, B., De Vlieger, D., and Saelens, X. (2018). Vaccine options for influenza: thinking small. Current opinion in immunology 53, 22–29.

26. Tao, W., Hurst, B.L., Shakya, A.K., Uddin, M.J., Ingrole, R.S., Hernandez-Sanabria, M., Arya, R.P., Bimler, L., Paust, S., Tarbet, E.B., et al. (2017). Consensus M2e peptide conjugated to gold nanoparticles confers protection against H1N1, H3N2 and H5N1 influenza A viruses. Antiviral research 141, 62–72.

27. Kim, K.H., Kwon, Y.M., Lee, Y.T., Kim, M.C., Hwang, H.S., Ko, E.J., Lee, Y., Choi, H.J., and Kang, S.M. (2018). Virus-Like Particles Are a Superior Platform for Presenting M2e Epitopes to Prime Humoral and Cellular Immunity against Influenza Virus. Vaccines 6.

28. Neirynck, S., Deroo, T., Saelens, X., Vanlandschoot, P., Jou, W.M., and Fiers, W. (1999). A universal influenza A vaccine based on the extracellular domain of the M2 protein. Nature medicine 5, 1157–1163.

29. Matić, S., Rinaldi, R., Masenga, V., and Noris, E. (2011). Efficient production of chimeric human papillomavirus 16 L1 protein bearing the M2e influenza epitope in Nicotiana benthamiana plants. BMC biotechnology 11, 106.

30. Petukhova, N.V., Gasanova, T.V., Stepanova, L.A., Rusova, O.A., Potapchuk, M.V., Korotkov, A.V., Skurat, E.V., Tsybalova, L.M., Kiselev, O.I., Ivanov, P.A., et al. (2013). Immunogenicity and protective efficacy of candidate universal influenza A nanovaccines produced in plants by Tobacco mosaic virus-based vectors. Current pharmaceutical design 19, 5587–5600.

31. Hashemi, H., Pouyanfard, S., Bandehpour, M., Noroozbabaei, Z., Kazemi, B., Saelens, X., and Mokhtari-Azad, T. (2012). Immunization with M2e-displaying T7 bacteriophage nanoparticles protects against influenza A virus challenge. PloS one 7, e45765.

32. Liu, W., Peng, Z., Liu, Z., Lu, Y., Ding, J., and Chen, Y.H. (2004). High epitope density in a single recombinant protein molecule of the extracellular domain of influenza A virus M2 protein significantly enhances protective immunity. Vaccine 23, 366–371.

33. Kim, M.C., Lee, J.S., Kwon, Y.M., O, E., Lee, Y.J., Choi, J.G., Wang, B.Z., Compans, R.W., and Kang, S.M. (2013). Multiple heterologous M2 extracellular domains presented on virus-like particles confer broader and stronger M2 immunity than live influenza A virus infection. Antiviral research 99, 328–335.

34. Cheng, W. (2016). The Density Code for the Development of a Vaccine? Journal of pharmaceutical sciences 105, 3223–3232.

35. Tao, P., Zhu, J., Mahalingam, M., Batra, H., and Rao, V.B. (2019). Bacteriophage T4 nanoparticles for vaccine delivery against infectious diseases. Advanced drug delivery reviews 145, 57–72.

36. Deng, L., Ibañez, L.I., Van den Bossche, V., Roose, K., Youssef, S.A., de Bruin, A., Fiers, W., and Saelens, X. (2015). Protection against Influenza A Virus Challenge with M2e-Displaying Filamentous Escherichia coli Phages. PloS one 10, e0126650.

37. Tao, P., Mahalingam, M., Marasa, B.S., Zhang, Z., Chopra, A.K., and Rao, V.B. (2013). In vitro and in vivo delivery of genes and proteins using the bacteriophage T4 DNA packaging machine. Proceedings of the National Academy of Sciences of the United States of America 110, 5846–5851.

38. Tao, P., Wu, X., Tang, W.C., Zhu, J., and Rao, V. (2017). Engineering of Bacteriophage T4 Genome Using CRISPR-Cas9. ACS synthetic biology 6, 1952–1961.

39. Shivachandra, S.B., Li, Q., Peachman, K.K., Matyas, G.R., Leppla, S.H., Alving, C.R., Rao, M., and Rao, V.B. (2007). Multicomponent anthrax toxin display and delivery using bacteriophage T4. Vaccine 25, 1225–1235.

40. Sathaliyawala, T., Rao, M., Maclean, D.M., Birx, D.L., Alving, C.R., and Rao, V.B. (2006). Assembly of human immunodeficiency virus (HIV) antigens on bacteriophage T4: a novel in vitro approach to construct multicomponent HIV vaccines. Journal of virology 80, 7688–7698.

41. Tao, P., Li, Q., Shivachandra, S.B., and Rao, V.B. (2017). Bacteriophage T4 as a Nanoparticle Platform to Display and Deliver Pathogen Antigens: Construction of an Effective Anthrax Vaccine. Methods Mol Biol 1581, 255–267.

42. Deloizy, C., Fossum, E., Barnier-Quer, C., Urien, C., Chrun, T., Duval, A., Codjovi, M., Bouguyon, E., Maisonnasse, P., Hervé, P.L., et al. (2017). The anti-influenza M2e antibody response is promoted by XCR1 targeting in pig skin. Scientific reports 7, 7639.

43. Sedova, E.S., Scherbinin, D.N., Lysenko, A.A., Alekseeva, S.V., Artemova, E.A., and Shmarov, M.M. (2019). Non-neutralizing Antibodies Directed at Conservative Influenza Antigens. Acta naturae 11, 22–32.

44. Boudreau, C.M., and Alter, G. (2019). Extra-Neutralizing FcR-Mediated Antibody Functions for a Universal Influenza Vaccine. Frontiers in immunology 10, 440.

45. Qi, M., Zhang, X.E., Sun, X., Zhang, X., Yao, Y., Liu, S., Chen, Z., Li, W., Zhang, Z., Chen, J., et al. (2018). Intranasal Nanovaccine Confers Homo- and Hetero-Subtypic Influenza Protection. Small (Weinheim an der Bergstrasse, Germany) 14, e1703207.

46. Saelens, X. (2019). The Role of Matrix Protein 2 Ectodomain in the Development of Universal Influenza Vaccines. The Journal of infectious diseases 219, S68–s74.

47. Eliasson, D.G., Omokanye, A., Schön, K., Wenzel, U.A., Bernasconi, V., Bemark, M., Kolpe, A., El Bakkouri, K., Ysenbaert, T., Deng, L., et al. (2018). M2e-tetramer-specific memory CD4 T cells are broadly protective against influenza infection. Mucosal immunology 11, 273–289.

48. Mezhenskaya, D., Isakova-Sivak, I., and Rudenko, L. (2019). M2e-based universal influenza vaccines: a historical overview and new approaches to development. J Biomed Sci 26, 76.

49. Deng, L., Roose, K., Job, E.R., De Rycke, R., Van Hamme, E., Gonçalves, A., Parthoens, E., Cicchelero, L., Sanders, N., Fiers, W., et al. (2017). Oral delivery of Escherichia coli persistently infected with M2e-displaying bacteriophages partially protects against influenza A virus. Journal of controlled release : official journal of the Controlled Release Society 264, 55–65.

50. Sun, X., Wang, Y., Dong, C., Hu, J., and Yang, L. (2015). High copy numbers and N terminal insertion position of influenza A M2E fused with hepatitis B core antigen enhanced immunogenicity. Bioscience trends 9, 221–227.

51. Gomes, A.C., Flace, A., Saudan, P., Zabel, F., Cabral-Miranda, G., Turabi, A.E., Manolova, V., and Bachmann, M.F. (2017). Adjusted Particle Size Eliminates the Need of Linkage of Antigen and Adjuvants for Appropriated T Cell Responses in Virus-Like Particle-Based Vaccines. Frontiers in immunology 8, 226.

52. Li, X., Xing, R., Xu, C., Liu, S., Qin, Y., Li, K., Yu, H., and Li, P. (2021). Immunostimulatory effect of chitosan and quaternary chitosan: A review of potential vaccine adjuvants. Carbohydrate polymers 264, 118050.

53. Berger, A. (2000). Th1 and Th2 responses: what are they? BMJ (Clinical research ed.) 321, 424.

54. Azouz, A., Razzaque, M.S., El-Hallak, M., and Taguchi, T. (2004). Immunoinflammatory responses and fibrogenesis. Medical electron microscopy : official journal of the Clinical Electron Microscopy Society of Japan 37, 141–148.

55. Von Holle, T.A., and Moody, M.A. (2019). Influenza and Antibody-Dependent Cellular Cytotoxicity. Frontiers in immunology 10, 1457.

56. Kim, M.C., Lee, Y.N., Ko, E.J., Lee, J.S., Kwon, Y.M., Hwang, H.S., Song, J.M., Song, B.M., Lee, Y.J., Choi, J.G., et al. (2014). Supplementation of influenza split vaccines with conserved M2 ectodomains overcomes strain specificity and provides long-term cross protection. Molecular therapy : the journal of the American Society of Gene Therapy 22, 1364–1374.

57. Tao, P., Mahalingam, M., and Rao, V.B. (2016). Highly Effective Soluble and Bacteriophage T4 Nanoparticle Plague Vaccines Against Yersinia pestis. Methods in molecular biology (Clifton, N.J.) 1403, 499–518.

